# Type I PRMT inhibitor MS023 promotes *SMN2* exon 7 inclusion and synergizes with nusinersen to rescue the phenotype of SMA mice

**DOI:** 10.1101/2022.10.18.512489

**Authors:** Anna J Kordala, Nina Ahlskog, Muhammad Hanifi, Amarjit Bhomra, Jessica Stoodley, Wooi Fang Lim, Suzan M Hammond, Matthew JA Wood, Carlo Rinaldi

**Author notes:** Address correspondence to: Carlo Rinaldi, Institute of Developmental and Regenerative medicine (IDRM), Old Road Campus, University of Oxford, OX3 7TY Oxford, United Kingdom.

## Abstract

Spinal muscular atrophy (SMA) is the leading genetic cause of infant mortality. The advent of approved treatments for this devastating condition has significantly changed SMA patients’ life expectancy and quality of life. Nevertheless, these are not without limitations, and research efforts are underway to develop new approaches to be used alone and in combination, to ensure improved and long-lasting benefits for SMA patients. Protein arginine methyltransferases (PRMT) are emerging as druggable epigenetic targets, with several small molecule PRMT inhibitors already in clinical trial stage. From a screen of highly potent and selective next generation epigenetic small molecules, we have identified MS023, a potent and selective type I PRMT inhibitor, able to promote *SMN2* exon 7 inclusion and increase SMN protein levels in preclinical SMA model, by inhibiting the binding of splicing factor hnRNPA1 to *SMN2* pre-mRNA. Treatment of SMA mice with MS023 results in amelioration of the disease phenotype, with strong synergistic amplification of the positive effect when delivered in combination with the *SMN2*-targeting antisense oligonucleotide nusinersen. Moreover, transcriptomic analysis revealed that MS023 treatment has very minimal off-target effects and that the added benefit of the combination therapy is mainly attributable to targeting neuroinflammation. Our study warrants further clinical investigation of PRMT inhibition both as a stand-alone and add-on therapy for SMA patients.

## Introduction

Spinal muscular atrophy (SMA) is a genetic neuromuscular condition, affecting 1:8,000– 10,000 live births (Pearn 1978) and, to this date, a leading inherited cause of infant mortality worldwide (Crawford and Pardo 1996). SMA is caused by inactivating mutations, mainly homozygous deletions, in the survival motor neuron 1 (*SMN1)* gene on chromosome 5 (Lefebvre et al. 1995). The encoded protein SMN is ubiquitously expressed, localizing both to the cytoplasmic and nuclear compartments within the cell, where it exerts numerous essential functions, including biogenesis of small nuclear ribonucleoproteins (snRNPs), its most widely studied function (Friesen, Massenet, et al. 2001; Pellizzoni, Yong, and Dreyfuss 2002; R. Zhang et al. 2011), 3’ processing of histone mRNAs (Tisdale et al. 2013), control of transcription (Strasswimmer et al. 1999; Suraweera et al. 2009; Yanling Zhao et al. 2016),

R-loop resolution (Suraweera et al. 2009; Yanling Zhao et al. 2016), RNA trafficking (Akten et al. 2011; Fallini et al. 2011, 2014, 2016; Hubers et al. 2011; Piazzon et al. 2008; Rage et al. 2013; Rossoll et al. 2003; Tadesse et al. 2008) and pre-mRNA splicing (Charroux et al. 1999; Makarov et al. 2012; Pellizzoni et al. 1998; Shafey et al. 2010). The *SMN2* gene is a centromeric copy of telomeric *SMN1*, with a critical C to T substitution in position 6 of exon 7, which creates an exonic splicing silencer site and is recognised by a splicing factor, hnRNPA1 (Kashima et al. 2007; Kashima and Manley 2003). As a consequence, the *SMN2* gene mainly encodes a shorter and rapidly degraded SMN isoform lacking exon 7 (Δ7 SMN), with only 10–15% of *SMN2* transcripts still capable of generating a full length SMN protein (Lorson et al. 1999; Monani et al. 1999). The number of *SMN2* copies varies in the general population and is the main modifier of disease severity identified so far, with higher number of copies being associated with a milder SMA phenotype (Feldkötter et al. 2002; McAndrew et al. 1997). Depending on the age of onset and motor milestones achieved, SMA has been divided into four clinical types (I–IV) (Munsat and Davies 1992), with type I SMA infants showing symptoms before 6 months of age and never gaining the ability to sit unaided.

Loss of SMN leads to degeneration of lower α-motor neurons by molecular mechanisms which are not fully understood. Other neuronal and non-neuronal cell populations are affected in SMA patients and include sensory neurons (Gogliotti et al. 2012; Jablonka et al. 2006; Martinez et al. 2012; Mentis et al. 2011; Rudnik-Schöneborn et al. 2003), skeletal muscle (Arnold et al. 2004; Kim et al. 2020; Martínez-Hernández et al. 2009) heart and vasculature (Araujo, Araujo, and Swoboda 2009; Bevan et al. 2010; Heier et al. 2010; Lipnick et al. 2019; Rudnik-Schöneborn et al. 2010; Somers et al. 2016), liver (Crawford et al. 1999; Deguise et al. 2019), and pancreas (Bowerman et al. 2012), supporting the notion that SMA is a multisystemic condition.

In the last decade, the advent of successful therapeutic approaches, combined with improvements in standard of care, have effectively changed the course of this disease, significantly slowing the progression of all SMA types (Harding et al. 2015; Eugenio Mercuri et al. 2020, 2022). One of such approved strategies entails intrathecal injections of nusinersen, an antisense oligonucleotide (ASO) targeting intronic splicing silencer N1 (ISS-N1), promoting *SMN2* exon 7 inclusion and increasing levels of full length SMN protein. Concomitantly, these treatment opportunities pose new challenges, including their yet-to-be determined long-term effects, the rise of new phenotypes in treated patients, which is particularly relevant for approaches such as nusinersen, solely targeting the central nervous system (CNS), the need for repeated invasive administrations, and high costs. Altogether, these considerations highlight the urgency for development of therapeutic combinations to address these important limitations and to provide additional benefit to patients. Epigenetic regulation of gene expression, which involves covalent and sequence-specific modifications of histone and non-histone proteins, is a dynamic and reversible process that establishes normal cellular phenotypes and, when dysregulated, contributes to a wide range of human diseases, including SMA (Allis et al. 2007; Cao et al. 2016; Hauke et al. 2009; Murray et al. 2015; Portela and Esteller 2010; Zheleznyakova et al. 2013). In recent years, key protein families that mediate epigenetic signalling through the acetylation and methylation of histones and non-histone proteins, including histone deacetylases (HDACs), protein methyltransferases (PRMT), histone lysine methyltransferases (KMTs) and demethylases (KDMs), and bromodomain-containing proteins (BRD) have emerged as attractive druggable targets using small molecules, due to the dynamic nature of disease-associated epigenetic states (Arrowsmith et al. 2012). Several small molecule inhibitors of histone deacetylases have been tested in SMA models (Mohseni, Zabidi-Hussin, and Sasongko 2013), but their lack of specificity, low potency, and poor understanding of their mechanisms of action have significantly limited their translation into the clinic (Kissel et al. 2014; Krosschell et al. 2018; E. Mercuri et al. 2007). A recent study has shown that nusinersen, while promoting exon 7 inclusion, also induces a silencing histone mark H3K9me2 on *SMN2* gene, creating a roadblock to RNA polymerase II elongation. The histone deacetylase inhibitor valproic acid counteracts the chromatin effects of the ASO, resulting in higher exon 7 inclusion upon combined treatment compared to nusinersen alone (Marasco et al. 2022). The primary aim of this project is to identify next generation small molecules targeting epigenetic proteins able to increase SMN protein and evaluate their therapeutic potential in SMA animal models alone and as an add-on treatment.

## Results

### Epigenetic screening of SMN2 modulators

In order to identify small molecules that selectively modulate *SMN2* pre-mRNA splicing to include exon 7, we performed a cell-based screen in SMA type II-patient derived fibroblasts carrying 3 copies of the *SMN2* gene, using a collection of 54 chemical probes from the Structural Genomics Consortium (SGC) collection (https://www.thesgc.org/chemical-probes) (Scheer et al. 2019; Wu et al. 2019) (Supplementary Table 1). This unique library includes compounds targeting key epigenetic regulatory proteins with a high degree of potency and selectivity, and a favourable therapeutic index (Ackloo, Brown, and Müller 2017). The maximum non-toxic concentrations for each compound, established by a viability assay in these cells, were used in the screen (Supplementary Figure 1). Of the 54 molecules, only one molecule, selective type I PRMT inhibitor MS023, was able to promote exon 7 inclusion in *SMN2* pre-mRNA, without affecting total *SMN2* mRNA levels (Figure 1A and B). MS023 treatment in these fibroblasts also increased SMN protein levels up to 1.6-fold, as determined by Western blot analysis (Figure 1C). Notably, treatment with PRMT5 inhibitors LLY-283 and GSK591 resulted in reduction of exon 7 inclusion in *SMN2* pre-mRNA and in SMN protein, respectively (Figure 1A–C), overall suggesting that protein arginine asymmetric and symmetric dimethylation by different families of PRMTs exert opposite effects on SMN regulation. Other compounds, including bromodomain inhibitors (BAY-299, BI-9564, JQ1) and lysine demethylase inhibitors (GSK-J1, GSK-LSD), also elicited a ≥1.5-fold increase in SMN protein without affecting mRNA levels, hinting at a direct or indirect effect on SMN protein regulation (Figure 1A-C). PRMTs are involved in several critical biological functions (Blanc and Richard 2017), and represent a promising therapeutic target for many human diseases from cancer to neurodegeneration, with at least eight PRMT inhibitors attaining clinical trial testing in human cancers (Guccione and Richard 2019; Hwang et al. 2021; Yang and Bedford 2013). Altogether, the direct effect of PRMT type I inhibition on *SMN2* exon 7 inclusion and the potential for clinical impact of this class of molecules, prompted us to further investigate MS023 as a therapeutic agent for SMA.

**Figure 1.**
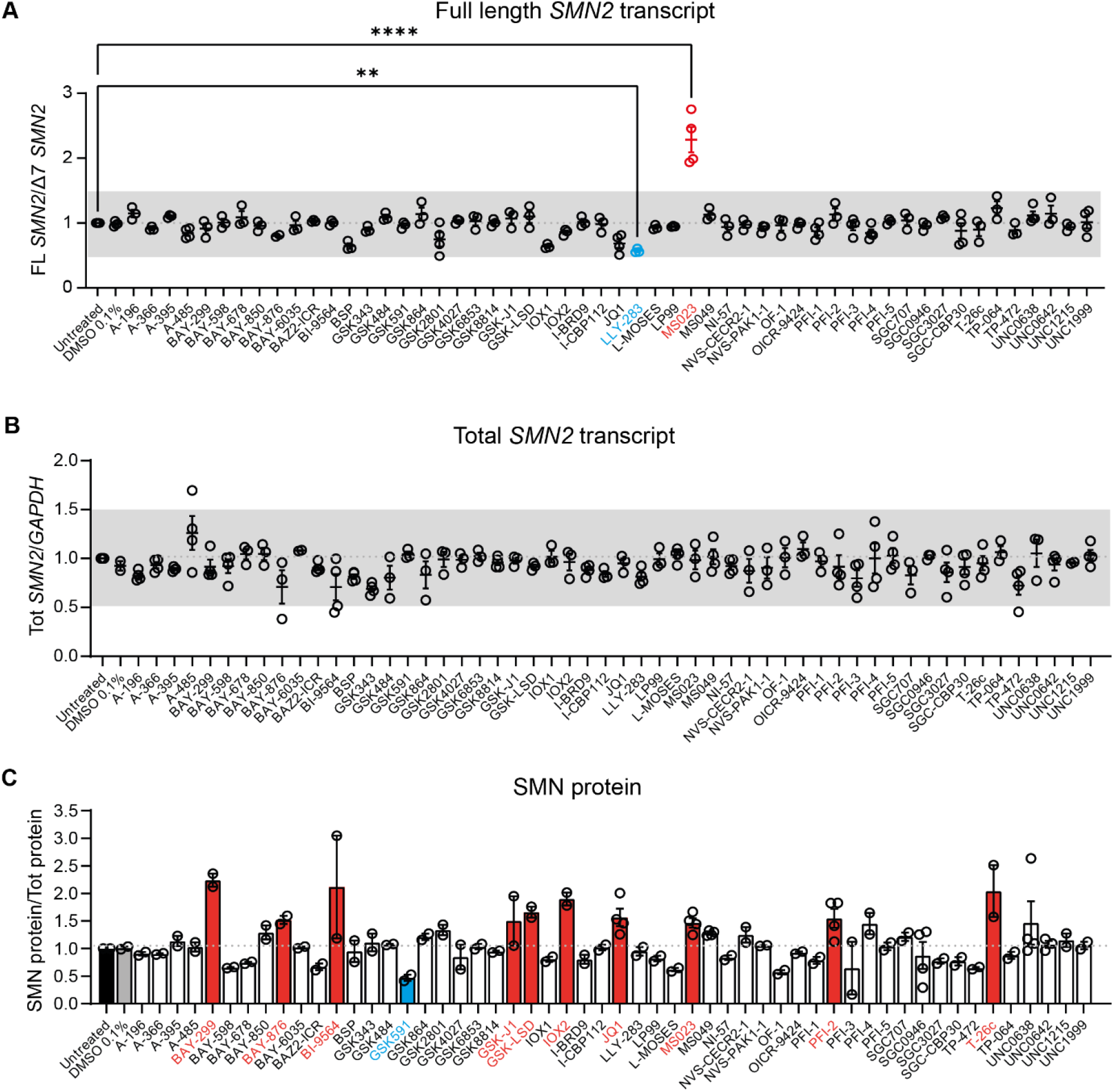
Screening of epigenetic small molecules in SMA fibroblasts. SMA type II patient-derived fibroblasts were treated with the indicated small molecule at the appropriate maximum tolerated dose (range: 1–10 μM) (*n* = 3–4). Cells were harvested for RNA (**A**–**B**) and protein (**C**) quantification 48 and 72 hours post treatment, respectively. **A**, Full length (FL) *SMN2* transcript levels relative to Δ7 *SMN*2 are expressed as fold change compared to untreated SMA fibroblasts, normalised to one (dashed line). FL *SMN2*/Δ7 *SMN2* ratios was significantly increased by MS023 and decreased by LLY-283 treatment. **B**, Tot *SMN2* transcript levels relative to *GAPDH* are expressed as fold change compared to untreated SMA fibroblasts, normalised to one (dashed line). **A**–**B**, Each dot represents a biological replicate (*n* = 3–4). Grey bar indicates values within the 0.5-1.5 range. **C**, SMN protein levels relative to total protein are expressed as fold change compared to untreated SMA fibroblasts, normalised to one (dashed line). Each dot represents a biological replicate (*n* = 2–4). Values ≥1.5 and ≤0.5 are depicted in red and blue, respectively. **A**-**C**, Data are represented as mean ± s.e.m. and compared with a one-way ANOVA test with multiple comparisons (\*\**P* ≤ 0.01; \*\*\*\**P* ≤ 0.0001).

### *Type I PRMT inhibition promotes exon 7 inclusion in* SMN2 *pre-mRNA by decreasing hnRNPA1 binding*

MS023 is a recently identified potent and selective inhibitor of type I PRMTs harbouring an ethylendiamino group, a critical moiety for its activity (Figure 2A) (Eram et al. 2016). Treatment of SMA fibroblasts with MS023 led to a dose-dependent increase in both *SMN2* exon 7 inclusion and protein levels (Figure 2B–D; Supplementary Figure 2A). This effect was not observed upon treatment with MS094 (Supplementary Figure 2B and C), an inactive MS023 analogue where the terminal primary amino group in the ethylenediamino group is replaced with a hydroxyl group (Eram et al. 2016), further confirming its dependency on PRMT activity. Knock-down of each and any combination of two of the type I PRMTs (PRMT 1, 3, 4, 6, and 8) resulted in no change in *SMN2* splicing, suggesting a high degree of functional overlap between these proteins (not shown). In order to identify the PRMT substrate mediating the effect of MS023 on *SMN2*, we interrogated a recently published dataset of the arginine methyl proteome in human NB4 cells upon treatment with MS023 (Fong et al. 2019). Out of 72 responsive targets, we noted that MS023 determined largely a downregulation of asymmetric dimethylarginine (ADMA) and an increase in monomethylarginine (MMA) sites in the heterogeneous nuclear ribonucleoprotein A1 (hnRNPA1) (Fong et al. 2019). hnRNPA1 directly binds *SMN2* pre-mRNA across multiple sites and is a well-established negative regulator of exon 7 splicing (Bose et al. 2008; Cartegni et al. 2006; Chen et al. 2008; Hua et al. 2008; Kashima, Rao, and Manley 2007; Koed Doktor et al. 2011; Singh et al. 2013; Xiao et al. 2012). Given that PRMT-regulated methylation regulates binding of RNA-binding proteins to RNAs (Blanc and Richard 2017), we hypothesised that MS023-induced ADMA-to-MMA switch in hnRNPA1 methylation affects its binding to *SMN*2 pre-mRNA, resulting in exon 7 inclusion. In order to test this hypothesis, we treated SMA fibroblasts with increasing concentrations of MS023 and confirmed a dose-responsive increase in both symmetric dimethylarginine (SDMA) and MMA methylation and a concomitant reduction in ADMA levels of hnRNPA1 protein (Figure 2E and F). Concomitantly, we observed a dose-responsive reduction of the fraction of FL *SMN2* transcripts bound to hnRNPA1, as determined by a crosslinking and immunoprecipitation (CLIP) assay (Figure 2G), which validated our working model (Figure 2H). Treatment with MS023 did not change hnRNPA1 levels nor its subcellular localization, further suggesting that the treatment-induced change in arginine methylation specifically affects hnRNPA1 binding affinity with its RNA target (Supplementary Figure 2D and E). Taken together, these findings indicate that MS023 promotes *SMN2* exon 7 inclusion via decreased binding of hnRNPA1.

**Figure 2.**
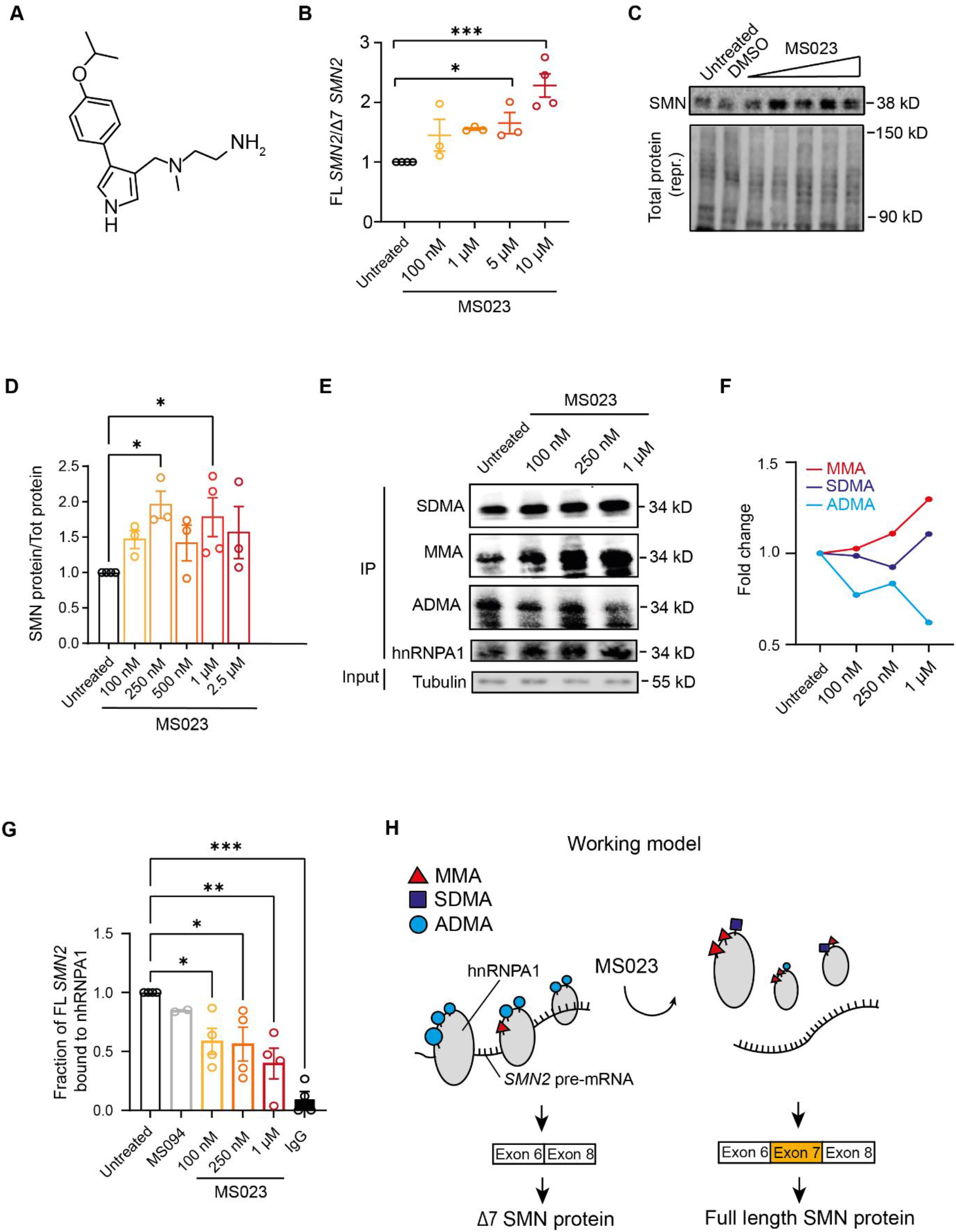
MS023 promotes exon 7 inclusion into *SMN2* transcript via hnRNPA1 binding inhibition. **A**, Chemical structure depiction of MS023 (PubChem CID: 92136227). **B-D** SMA type II patient-derived fibroblasts were treated with the indicated concentration of MS023 (range: 100 nm–10 μM), (*n* = 3–4). Cells were harvested for RNA (**B**) and protein (**C, D**) quantification 48 and 72 hours post treatment, respectively, **B**. Full length (FL) *SMN2* transcript levels relative to Δ7 *SMN*2 are expressed as fold change compared to untreated SMA fibroblasts, normalised to one. Each dot represents a biological replicate (*n* = 3–4). **C**, Western blot showing SMN protein levels upon treatment with increasing MS023 concentrations (top). A representative section of total protein stain, used for protein normalization, is shown (bottom). Size in kilodalton is indicated on the right. **D**, Quantification of SMN protein levels relative to total protein is shown. Each dot represents a biological replicate (*n* = 3). **E**, Arginine monomethylation (MMA) and symmetric dimethylation (SDMA) of hnRNPA1 are increased and asymmetric dimethylation (ADMA) of hnRNPA1 is decreased upon MS023 treatment, as shown by hnRNPA1 immunoprecipitation followed by a Western blot. SMA type II patient-derived fibroblasts were treated with the indicated concentration of MS023 (range: 100 nm–1 μM). hnRNPA1 was immunoprecipitated and Western blots were performed using and anti-MMA, anti-SDMA, anti-ADMA, anti-hnRNPA1 and anti-tubulin antibodies. **F**, Quantification of arginine methylation changes normalized to total hnRNPA1 is shown. **G**, Ratio of FL *SMN2* transcripts bound to hnRNPA1 protein and relative to *GAPDH* was assayed using crosslinking immunoprecipitation (CLIP) in SMA patient fibroblasts treated with increasing concentration of MS023. IgG antibody and MS094 were used controls, MS094 is a negative control for MS023 lacking PRMT inhibition property. Each dot represents a biological replicate (*n* = 2-4). **B, D, G** Data are represented as mean ± s.e.m. and compared with a one-way ANOVA test with multiple comparisons (\**P* ≤ 0.05; \*\**P* ≤ 0.01; \*\*\**P* ≤ 0.001). **H**, Representation of the mechanism of exon 7 inclusion into the *SMN2* transcript by MS023. Blue circles, purple squares and red triangles represent asymmetric dimethylarginine (ADMA), symmetric dimethylarginine (SDMA) monomethylarginine (MMA) respectively.

### Oral administration of MS023 improves the phenotype of SMA mice alone and in synergy with nusinersen

We next evaluated the efficacy and tolerability of MS023 treatment *in vivo* in a severe preclinical mouse model of SMA (Martínez-HernáLi et al. 2000). These mice, which lack the mouse *Smn* gene and only carry a single copy of the human *SMN2* gene, display a phenotype with weight loss and reduced motor activity starting at postnatal day 5 (P5) and typically reach humane end point by P9. Daily oral administration of MS023 or vehicle (0.5% DMSO in 0.9% saline solution) was performed in SMA mice from P0 until reaching a humane end point (Figure 3A). We tested a range of doses (1, 2, 5, and 40 mg/kg) and found that treatment with both 2 mg/kg and 5 mg/kg resulted in significant increase in survival, with the 2 mg/kg dose achieving the best effect (median: 10 days), compared to vehicle-treated mice (median: 6 days; *P* < 0.0001) (Figure 3B). Mice treated with this regimen also showed an improvement in the disease-associated weight loss (Figure 3C). No further amelioration was observed with 40 mg/kg MS023, hinting that with this dose the therapeutic window has been surpassed (Figure 3B, Supplementary Figure 3A). Notably, we detected an increase in full length (FL) *SMN2* transcript and SMN protein levels in the tissues mostly affected in the disease, spinal cord and skeletal muscle (Eugenio Mercuri et al. 2022), of SMA mice treated with 2 mg/kg MS023 (Figure 3D–F; Supplementary Figure 3B). Since the mechanism of action of the 2’-O-methoxyethyl phosphorothioate-modified drug nusinersen, a currently approved ASO therapy for SMA patients, consists of promoting exon 7 inclusion into *SMN2* pre-mRNA by blocking the recruitment of hnRNP splicing repressors at the ISS-N1 site (Chiriboga et al. 2016; Finkel et al. 2016, 2017; Haché et al. 2016; Eugenio Mercuri et al. 2018), we postulated that a combinatorial treatment with MS023 would synergistically lead to improved therapeutic benefit and allow a cost-effective ASO dosing regimen in SMA. In order to test this hypothesis, SMA mice were treated at P0 with a single subcutaneous administration of 30 mg/kg nusinersen, a minimum effective dose previously shown to extend the SMA mice life span (Hammond et al. 2016), alone or in combination with daily oral administration of 2 mg/kg MS023 from P1 to P6 (Figure 4A).

**Figure 3.**
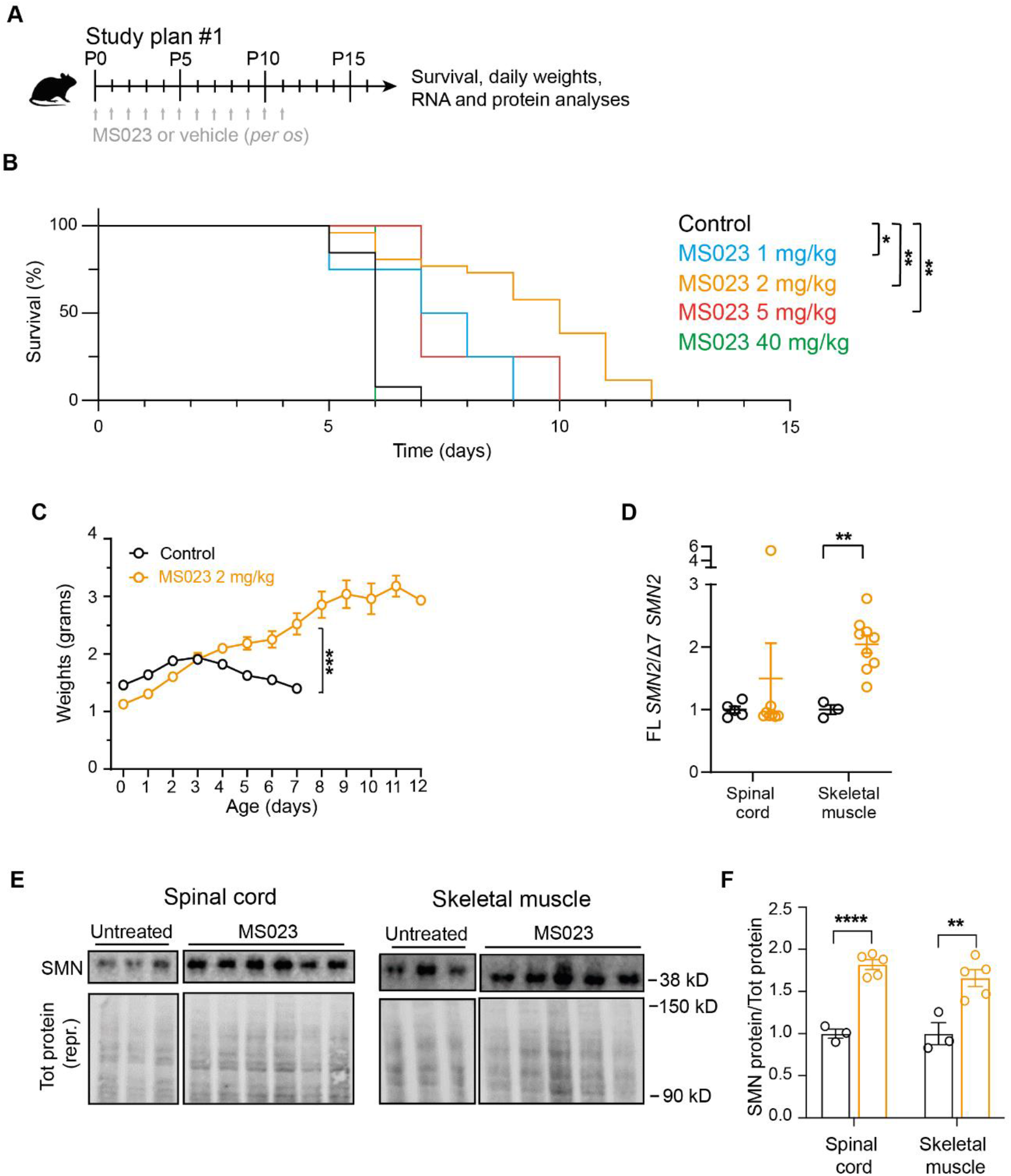
Oral administration of MS023 improves the phenotype of SMA mice. **A**, Diagram of the study design; mice were treated daily with oral administrations of MS023 or vehicle (0.5% DMSO in saline) from postnatal day 0 (P0) using a Hamilton syringe. **B**, Kaplan-Meier survival estimation of mice treated with vehicle (*n* = 14), 1 mg/kg MS023 (*n* = 4), 2 mg/kg MS023 (*n* = 26), 5 mg/kg MS023 (*n* = 4), or 40 mg/kg MS023 (*n* = 4) SMA mice. **C**, Body weights of MS023- and vehicle-treated SMA mice from postnatal day 0 are shown. **D**, FL *SMN2* transcript levels relative to Δ7 *SMN2* in spinal cord and skeletal muscle of treated mice compared to vehicle-treated mice, normalised to one. Each dot represents a biological replicate (*n* = 3–9). **E**, Western blot showing SMN protein levels upon MS023 treatment in spinal cords and skeletal muscles of treated mice (top). A representative section of total protein stain, used for protein normalization, is also shown (bottom). Size in kilodalton is indicated on the right. **F**, Quantification of SMN protein levels in the spinal cord and skeletal muscle relative to total protein is shown. Each dot represents a biological replicate (*n* = 3–8). **B–D**, and **F**, Data are represented as mean ± s.e.m. and compared with Mantel-Cox test (\**P* ≤ 0.05; \*\**P* ≤ 0.01) (**B**) or a one-way ANOVA test with multiple comparisons (\*\**P* ≤ 0.01; \*\*\**P* ≤ 0.001, \*\*\*\**P* ≤ 0.0001) (**C, D, F**).

**Figure 4.**
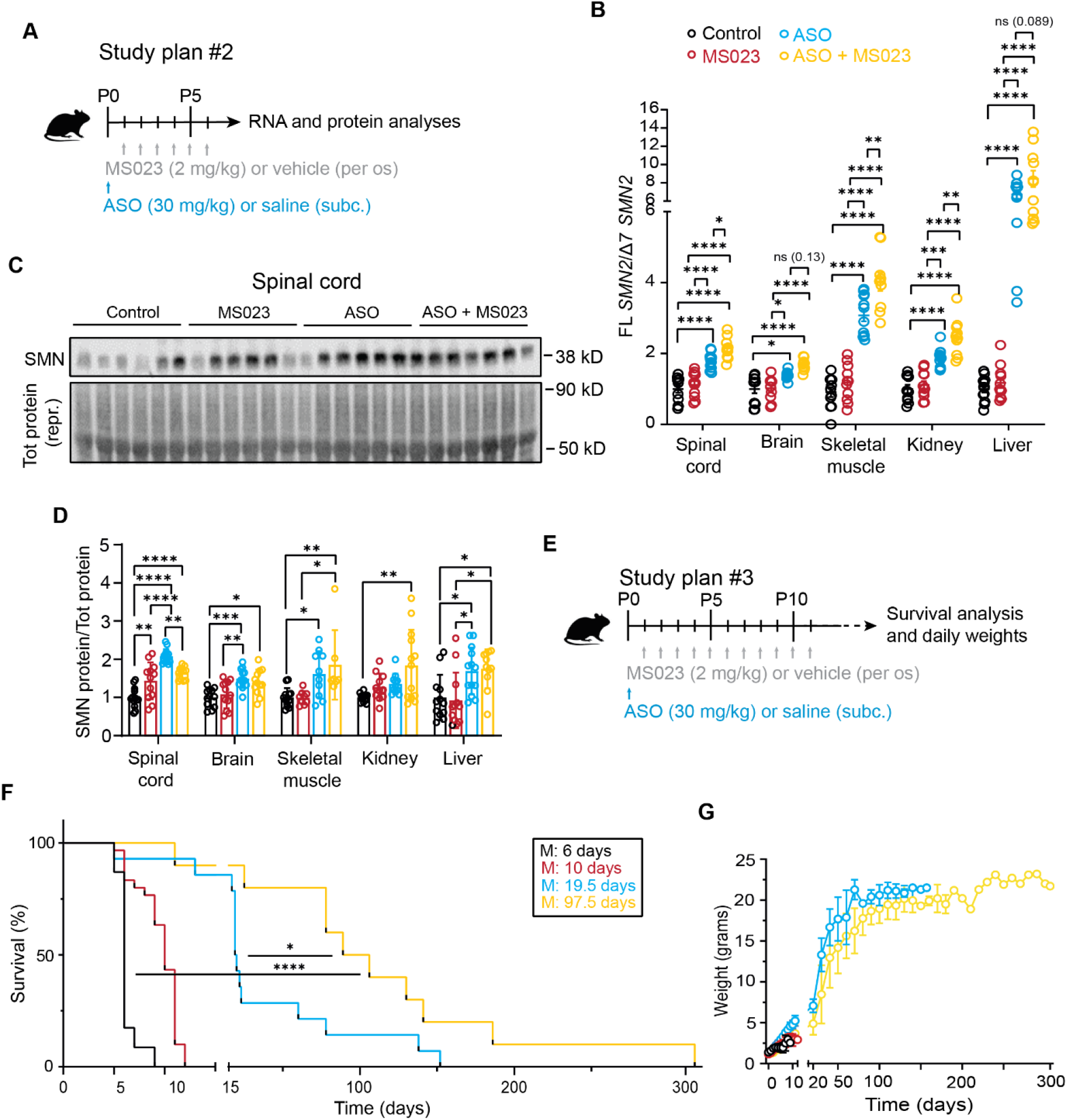
Combinatorial treatment with MS023 and ASO exerts synergistic effects in SMA mice. **A**, Diagrams of the study design. Study plan #2: At P0 SMA mice were injected subcutaneously with *SMN2*-targeting ASO (30 mg/kg) or saline; from P1 mice were treated daily with oral administrations of MS023 (2 m/kg) or vehicle (0.5% DMSO in saline) using a Hamilton syringe until P6. At P7, mice were sacrificed and tissues collected for analysis. **B**, FL *SMN2* transcript levels relative to Δ7 *SMN2* in spinal cord, brain, skeletal muscle, kidney and liver of treated SMA mice compared to vehicle-treated SMA mice, normalised to one. Each dot represents a biological replicate (*n* = 10–12). **C**, Western blot showing SMN protein levels upon the indicated treatments in spinal cords of SMA mice (top). A representative section of total protein stain, used for protein normalization, is also shown (bottom). Size in kilodalton is indicated on the right. **D**, Quantification of SMN protein levels relative to total protein is shown. Each dot represents a biological replicate (*n* = 7–13). **E**, Study design #3: At P0 mice were injected subcutaneously with *SMN2*-targeting ASO (30 mg/kg) or saline. From P1 mice were treated daily with oral administrations of MS023 (2 m/kg) or vehicle (0.5% DMSO in saline) using a Hamilton syringe until P12. Mice were weighed daily until they reached their humane end point. **F**, Kaplan-Meier survival estimation of SMA mice treated with *SMN2*-targeting ASO (*n* = 15), MS023 (*n* = 14), *SMN2*-targeting ASO and MS023 (*n* = 10), and vehicle (*n* = 23). **G**, Body weights of SMA mice from postnatal day 0 are shown. **B, D, F**, and **G**, Data are represented as mean ± s.e.m. and compared with a one-way ANOVA test with multiple comparisons (\**P* ≤ 0.05; \*\**P* ≤ 0.01; \*\*\**P* ≤ 0.001, \*\*\*\**P* ≤ 0.0001) or (**F**) Mantel-Cox test (\**P* ≤ 0.05; \*\*\*\**P* ≤ 0.0001).

Analysis of tissues collected at P7 showed that the combinatorial treatment was able to further enhance exon 7 inclusion in *SMN2* pre-mRNA (Figure 4B) and increase SMN protein levels (Figure 4C and D; Supplementary Figure 4), both in CNS and peripheral tissues. A further follow-up study designed to assess the effect on survival (Figure 4E) revealed that the combinatorial treatment of nusinersen and MS023 dramatically prolonged the lifespan (median: 97.5 days) compared to nusinersen alone (median: 19.5; *P* = 0.02) (Figure 4F), and improved the body weight of SMA mice (Figure 4G). Overall, these results suggest that oral administration of MS023 synergises with nusinersen to provide a therapeutic benefit in SMA.

### Molecular signature of the combinatorial treatment

In order to understand the molecular underpinnings of the added benefit provided by the combinatorial treatment, we performed bulk transcriptomic analysis in spinal cords of symptomatic (P7) SMA mice treated with MS023 only, nusinersen only, and MS023 in combination with nusinersen, compared to wild type and untreated SMA mice, following the treatment paradigm described above (Figure 4A). Out of a total of 5509 significantly dysregulated transcripts in SMA (*P* < 0.05, false discovery rate < 0.05), 2482/2874 (86%) of the upregulated and 2060/2635 (83%) of the downregulated genes were commonly corrected by nusinersen treatment alone and by combinatorial treatment (Figure 5A). Combinatorial treatment was able to exclusively rescue 359 (6.5% out of 5509) additional genes that were not corrected upon treatment with nusinersen alone (Supplementary Table 2). We generated a heatmap of the top significant genes (*P* < 0.01) of this category (Figure 5B) and performed a hallmark analysis, to depict and identify the changes that could explain the beneficial effects of the combinatorial treatment (Figure 5C and D). TNF-α signalling, together with several other immune-related pathways, including interferon response and complement activation, were highly enriched, overall suggesting that targeting neuroinflammation for therapy is key to achieving a beneficial effect in SMA. Interestingly, astrocyte dysfunction and chronic microglia activation have been observed early in SMA and other neurological conditions (Eikelenboom et al. 2002; Heneka, Kummer, and Latz 2014; Sargsyan, Monk, and Shaw 2005; Vukojicic et al. 2019) and their contribution to neuronal dysfunction and death is abundantly supported (Martin et al. 2017; Mcgivern et al. 2013; Rindt et al. 2015; Zhou, Feng, and Ko 2016). Given the role of SMN in RNA biogenesis and spliceosomal proteins assembly (Pellizzoni et al. 2001), and the observation of increased mis-splicing events upon SMN depletion (Bäumer et al. 2009; Custer et al. 2016; Doktor et al. 2017; Huo et al. 2014; Z. Zhang et al. 2008, 2013) we wondered whether the beneficial effect of the combinatorial treatment was due to restoration in the splicing profile. We identified 446 mis-splicing events in the spinal cord of SMA mice, with the vast majority (*n* = 353, 79%) being skipped exons, in accordance with previous reports (Bäumer et al. 2009; Custer et al. 2016; Doktor et al. 2017; Huo et al. 2014; Z. Zhang et al. 2008, 2013) (Supplementary Figure 5A and B). Individual treatment with MS023 or nusinersen were both able to restore the splicing profile almost fully (347/446 and 368/446, respectively), with only 14 unique splicing events corrected exclusively upon the combinatorial treatment (Figure 5E; Supplementary Table 3), suggesting that mis-splicing has a low threshold for normalization in SMA and is a poor predictor of treatment response.

**Figure 5.**
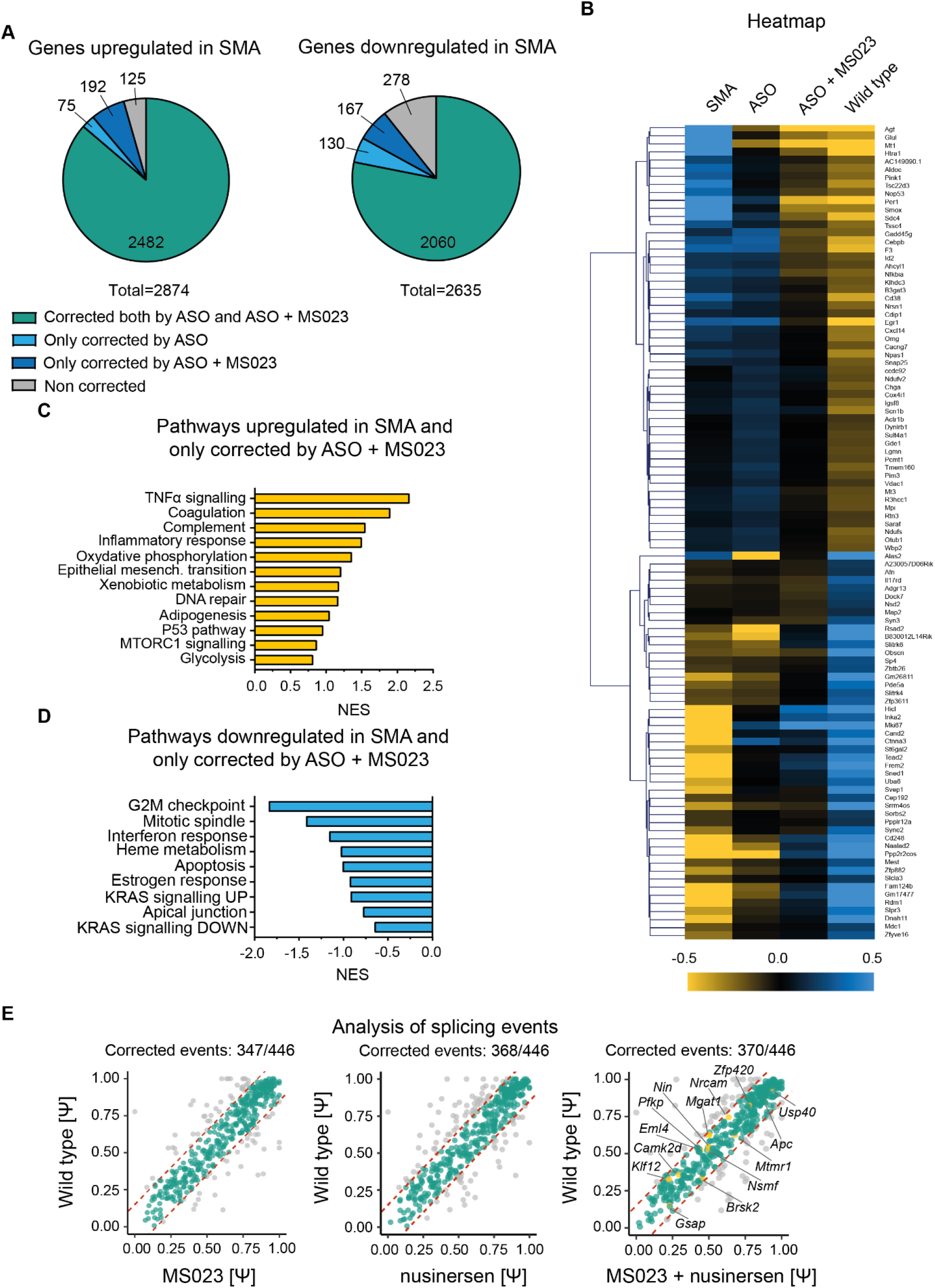
Combinatorial treatment with MS023 and ASO results in improved correction of the SMA transcriptomic signature compared to ASO alone. **A**, Pie charts showing the proportion of transcripts normalised upon the treatments, among the genes that are upregulated (left) and downregulated (right) in SMA mice. **B**, Normalised counts were used to generate the hierarchical clustering heatmap. Upregulated and downregulated genes are displayed in yellow and blue, respectively. **C**, Hallmark pathway analysis of genes upregulated and **D**, downregulated in SMA corrected by the combinatorial treatment only. **E**, Plot charts show the distribution of splicing events (Ψ) in SMA mice upon treatment relative to wild type littermates. The red dotted lines mark the ± 15% normalization range, corrected values are indicated in green.

## Discussion

Thousands of SMA patients are treated each year with nusinersen, the first SMA treatment to receive FDA and EMA approval in 2016 and 2017, respectively. Despite having enormously changed the life expectancy and quality of life of patients, this approach is not without limitations: it fails to address the peripheral manifestations of the disease, is costly, and the intrathecal administrations require patients’ hospitalization and well-trained clinicians, becoming more troublesome with time as the disease progresses (Eugenio Mercuri et al. 2020). Research efforts are underway to develop both new SMN-dependent and SMN-independent therapies for use in combination with current therapies, to provide improved and sustained benefit for all SMA patients.

From a screen of potent and highly selective epigenetic probes, we have identified MS023, a potent and selective next-generation type I PRMT inhibitor (Eram et al. 2016), able to elevate SMN protein in SMA models, at levels comparable to recently clinically approved small molecule risdiplam (Naryshkin et al. 2014) and to synergistically amplify the effects of nusinersen. Interestingly, type II PRMT-targeting by selective PRMT5 inhibitors GSK591 and LLY-283 reduced FL *SMN2*, supporting a mechanism of regulation of *SMN2* by this class of enzymes.

Protein arginine methylation is a prevalent posttranscriptional modification occurring at the nitrogen of guanidinium group of over 4000 proteins, many of which are RNA-binding proteins, influencing their stability, function, and interaction with other macromolecules (Hornbeck et al. 2015). In mammals, arginine methylation of histone and non-histone proteins is performed by nine PRMT enzymes, divided into three families: type I PRMTs (PRMT1, PRMT2, PRMT3, PRMT4, PRMT6 and PRMT8) perform monomethylation followed by an asymmetric dimethylation (ADMA) of arginine, with both methyl groups on a single guanidino nitrogen atom; type II PRMTs (PRMT5 and PRMT9) catalyse monomethylation, followed by a symmetric dimethylation (SDMA) where one methyl group is transferred to each of the nitrogen atoms; and type III PRMT (PRMT7) only performs monomethylation (MMA) (Guccione and Richard 2019). PRMTs have been shown to compete over the same substrates, frequently with opposite functional effects (Dhar et al. 2013).

Several studies have highlighted a link between PRMTs and SMN: To ensure fidelity of loading onto the correct snRNAs, three of the core spliceosomal Sm proteins (SmB, SmD1 and SmD3) undergo symmetric dimethylation by PRMT5, a modification recognised by the Tudor domain of SMN (Friesen, Paushkin, et al. 2001; Meister et al. 2001). SMN interaction with senataxin, a DNA-RNA helicase, depends on the dimethylated arginine and is reduced upon PRMT5 knockdown (Yanling Zhao et al. 2016). Lastly, levels of PRMT4 (CARM1) are upregulated in SMA mice spinal cord and patients’ cells (Sanchez et al. 2015). PRMTs represent a promising therapeutic target for many human diseases from cancer (Drew et al. 2017; Wang et al. 2016; Wu et al. 2022) to neurodegeneration (Dormann et al. 2012; Scaramuzzino et al. 2015; Suárez-Calvet et al. 2016), with eight PRMT inhibitors attaining clinical trial testing in human cancers (Blanc and Richard 2017; Guccione and Richard 2019; Hwang et al. 2021; Yang and Bedford 2013). Consistently with previous methylome analyses in cells treated with type I PRMT inhibitors, GSK3368712 (Noto et al. 2020) or MS023 (Karuppagounder et al. 2016), we propose that MS023 mechanism of action entails a switch in the arginine methylation profile from ADMA to SDMA/MMA of hnRNPA1, a major negative regulator of *SMN2* exon 7 inclusion (Bose et al. 2008; Cartegni et al. 2006; Chen et al. 2008; Hua et al. 2008; Kashima, Rao, and Manley 2007; Koed Doktor et al. 2011; Singh et al. 2013; Xiao et al. 2012), resulting in reduced binding affinity to *SMN2* pre-mRNA. Notably, nusinersen also acts by interfering with hnRNPA1 binding to the ISS-N1 sequence, suggesting a synergistic convergence on this mechanism of regulation. Since splicing is regulated by several RNA binding proteins, we cannot exclude that methylation of splicing factors other than hnRNPA1 also contribute, directly or indirectly, to the effect of MS023 on *SMN2* (Ravindra N. Singh et al. 2018).

Hallmark analysis of the 359 genes whose expression levels in spinal cords were only corrected by the combinatorial treatment, revealed enrichment of immune-related pathways, such as TNF-α signalling, interferon response and complement activation, overall suggesting that therapeutic targeting of neuroinflammation is key to achieving optimal and long-lasting effect in SMA. Interestingly, astrocyte dysfunction and chronic microglia activation have been observed early in SMA and other neurological conditions (Eikelenboom et al. 2002; Heneka, Kummer, and Latz 2014; Sargsyan, Monk, and Shaw 2005; Vukojicic et al. 2019) and play a determinative role in the disease pathogenesis (Martin et al. 2017; Mcgivern et al. 2013; Rindt et al. 2015; Zhou, Feng, and Ko 2016). Transcriptomic data in the preclinical model indicates that, at the dose employed in this study, MS023 shows a favourable safety profile, with minimal off-target effects both on gene expression and splicing alterations, in stark contrast to the pleiotropic effects of other epigenetic modulators such as valproic acid. Altogether, these promising preclinical results warrant further clinical investigations of MS023 or other selective type I PRMT inhibitors both as a stand-alone and an add-on treatment with nusinersen in SMA patients.

## Methods

### Small molecules screening

The library of small molecules was obtained from the Structural Genomic Consortium (SGC) (Williamoson AR, 2000). Upon reception the molecules were diluted in DMSO to 10 mM, aliquoted and stored at −20°C for the duration of the study. All the compounds, their targets and doses used in the study are listed in Supplementary Table 1.

### Cell lines and culture

Cells were grown in a humidified incubator at 37°C with 5% CO_2_. SMA type II patient fibroblasts were obtained from Coriell Institute (GM03813). The cells were maintained in Dulbecco’s Modified Eagle Medium GlutaMAX (Gibco) supplemented with 10% foetal bovine serum (Gibco) and 1× Antibiotic-Antimycotic (Gibco). SMA patient fibroblasts were plated in triplicates, in 12-well plates at 50,000 cells per well in 500 μL of medium for RNA or in duplicates in 6-well plates at 100,000 cells per well in 1 mL of medium for protein. After 6 h, compounds of interest were diluted in medium to 2× final concentration and added to the cells. RNA was isolated after 48 h incubation, protein after 72 h.

### Cell viability assay

MTS Cell Proliferation Assay Kit (Abcam) was used to determine the highest non-toxic concentration of small molecules in SMA patient fibroblasts. Briefly, 3000 cells/wells were plated in triplicates in a 96-well plate in half of the final volume of medium (100 μL). After 6 h, compounds of interest were diluted to 2× final concentration in 100 μL of medium and added in a range of concentrations (1 μM to 10 μM). 1 μM of staurosporine (Sigma) was used as a positive control for cell death. After 48 h, 20 μL of MTS reagent was added to each well. Four hours later, the absorbance was measured at 490 nm on CLARIOstar plate reader (BMG Labtech). The concentration was deemed non-toxic if the absorbance was not significantly different from the control and cells looked viable after visual inspection. In subsequent experiments, only the highest non-toxic concentration of each compound was used, unless stated otherwise.

### RNA/cDNA preparation and RT-qPCR

RNA extraction was performed using Maxwell® RSC simply RNA Kit (Promega) following the manufacturer’s instructions. The concentration was measured using Nanodrop 1000 spectrophotometer (Thermo Fisher), and cDNA was generated using an ABI High Capacity cDNA Reverse Transcription Kit (Invitrogen) following the manufacturer’s instructions. A qPCR reaction using Power SYBR Green Master Mix (Life Technologies) was performed and analysed on an Applied Biosystems StepOnePlus™ real-time PCR system (Life Technologies). FL *SMN2*, Δ7 *SMN2*, Tot *SMN2, PolJ, Gapdh* and *GAPDH* transcripts were amplified using gene- and species-specific primers (Supplementary Table 4).

### Protein extraction and Western blot

Proteins were harvested from ∼30 mg of tissue (*in vivo*) or two 6-well plates (*in vitro*) and homogenised in Ripa buffer with complete mini-proteinase inhibitors (Roche). Preparation of nuclear and cytoplasmic extracts from human fibroblasts was performed using a NE-PER Nuclear and Cytoplasmic Extraction Reagents (Thermo Fisher), as per manufacturer’s instructions. Proteins (10–15 μg from cells and 20–30 μg from tissue) were probed for human SMN protein using anti-SMN, clone SMN-KH monoclonal IgG1 (Sigma, MABE230), Histone 3 (Cell Signalling, 9715), Vinculin (Sigma, 062M4762), hnRNPA1 (Santa Cruz, 4B10), MMA (Cell Signalling, 8015), ADMA (Sigma Aldrich, 07-414), SDMA (Cell Signalling, 13222), β-tubulin (Abcam, 108342) and FAST green total protein stain and secondary antibody IRDye 800CW goat anti-mouse IgG (LI-COR Biosciences). Membranes were imaged on a LI-COR Odyssey FC imager and analysed with Image Studio™ software (LI-COR Biosciences).

### Crosslinking and immunoprecipitation assay (CLIP)

Briefly, for each condition, four 150 cm^2^ cell dishes of human patient fibroblasts were treated with 100 nM, 250 nM or 1 μM of MS023 for 48 h. After the incubation, the medium was aspirated and the cells were subjected to 150 mJ/cm^2^ 254 nm UV light in a Stratalinker UV Crosslinker and pelleted. The cells were then lysed in NP-40 lysis buffer and hnRNPA1 immunoprecipitation was performed with Dynabeads Protein G (Thermofisher) and hnRNAP1 antibody (Santa Cruz). Following DNA and protein digestion, RNA was isolated and cDNA was generated as previously outlined. A qPCR reaction using Power SYBR Green Master Mix (Life Technologies) was performed and analysed on an Applied Biosystems StepOnePlus™ real-time PCR system (Life Technologies). FL *SMN2* and *GAPDH* transcripts were amplified using gene- and species-specific primers.

### Protein immunoprecipitation

To detect changes in hnRNPA1 arginine methylation, SMA patient fibroblasts were either left untreated or treated with increasing concentrations MS023 (100 nM, 250 nM or 1 μM) for 48h. Cell pellets were lysed in NP-40 buffer and hnRNPA1 immunoprecipitation was performed with Dynabeads Protein G (Thermofisher, 10003D) and hnRNAP1 antibody (Santa Cruz), following the manufacturer’s protocol. Subsequently, Western blots were run according to the protocol described above.

### Mice

Mice were housed and all the procedures were carried out at the Biomedical Services Building, University of Oxford and authorised by the UK Home Office in accordance with the Animals (Scientific Procedures) Act 1986 and by the University of Oxford ethics committee (PPL no: PDFEDC6F0). All experiments were performed on the SMA mouse strain FVB.Cg-Smn1^tm1Hung^Tg(SMN2)2Hung/J – the ‘Taiwanese’ model (*Smn*^*−*/*−*^;*hSMN2*^+/*−*^), generated and maintained as previously described (Gogliotti et al. 2010; Hsieh-Li et al. 2000). MS023 was administered daily orally from P0 or P1 using a Hamilton syringe (Hamilton). Doses of 0, 1, 2, 5 or 40 mg/kg of MS023 were diluted in 0.5% DMSO and 0.9% saline and administered at a volume of 5 μL/g of body weight. Nusinersen (sequence: U*sC*sA*sC*sU*sU*sU*sC*sA*sU*sA*sA*sU*sG*sC*sU*sG*sG*s, where ‘S’ is phosphorothioate backbone and ‘*’ is a 2’-O-(2-Methoxyethyl)-oligoribonucleotides chemistry) was diluted in 0.9% saline and given once at 20 μL/g body weight, via subcutaneous injection at P0, in a dose of 30 mg/kg. Weights were recorded and overall health was assessed daily. Since the pups were treated daily from P0 and identifying and marking individual pups between P0 and P7 poses a high risk of misidentifying an animal, in this study a litter constitutes an experimental unit and all mice in the litter were subjected to the same treatment. Only litters between 7 and 11 pups were used to correct for average weight (mice in smaller litters tend to be bigger and live longer and the opposite is true for litter of 12 and above) and treatment was allocated randomly to a litter before it was born. All the experimental units treated were included in the analysis. Personnel performing daily weights and welfare checks for combinatorial therapy were blinded, researcher performing oral administration, injections and data analysis was not blinded. For survival analysis, the humane end point was reached upon 15% weight loss from a maximum weight or when the mouse was not able to right itself for 30 s. Mice were culled by decapitation (if younger than postnatal day 10) or cervical dislocation (if 10 days old or older). Tissues were harvested at the indicated postnatal day.

### RNA sequencing

Transcriptomic analysis was performed by Novogene (UK) company Limited (https://en.novogene.com/) on P7 spinal cords of mice in the following treatment groups: untreated SMA mice, SMA mice treated with nusinersen, SMA mice treated with MS023, SMA mice treated with nusinersen and MS023, untreated controls (4 biological replicates in each group). RNA quantification and integrity were assessed using the RNA Nano 6000 Assay Kit of the Bioanalyzer 2100 system (Agilent Technologies, CA, USA). mRNA was purified using poly-T oligo-attached magnetic beads. Fragmentation was carried out using divalent cations under elevated temperature in First Strand Synthesis Reaction Buffer. First strand cDNA was synthesised using random hexamer primer and M-MuLV Reverse Transcriptase. Second strand cDNA synthesis was subsequently performed using DNA Polymerase I and RNase H. Remaining overhangs were converted into blunt ends via exonuclease/polymerase activities. After adenylation of 3’ ends of DNA fragments, adaptor with hairpin loop structure were ligated to prepare for hybridization. In order to select cDNA fragments of preferentially 370∼420 bp in length, the library fragments were purified with AMPure XP system (Beckman Coulter, Beverly, USA). PCR products were purified (AMPure XP system) and library quality was assessed on the Agilent Bioanalyzer 2100 system. The clustering of the index-coded samples was performed on a cBot Cluster Generation System using TruSeq PE Cluster Kit v3-cBot-HS (Illumina) according to the manufacturer’s instructions. After cluster generation, the library preparations were sequenced on an Illumina Novaseq platform with a coverage of 25 million reads and 150 bp paired-end reads were generated. Raw data (raw reads) of fastq format were firstly processed through in-house perl scripts. In this step, clean data (clean reads) were obtained by removing reads containing adapter, reads 1 containing ploy-N and low-quality reads from raw data. Mus Musculus (GRCm38/mm10) reference genome was used, index of the reference genome was built using Hisat2 v2.0.5 and paired-end clean reads were aligned to the reference genome using Hisat2 v2.0.5. The mapped reads of each sample were assembled by StringTie (v1.3.3b) (Pertea et al. 2015) in a reference-based approach. Quantification of gene expression level Feature Counts v1.5.0-p3 was used to count the reads numbers mapped to each gene. Differential expression analysis was performed using the DESeq2 R package (1.20.0), Benjamini-Hochberg-adjusted *P*-values reported. Corrected *P*-value of 0.05 and absolute fold change of 2 were set as the threshold for significantly differential expression. Alternative splicing analysis rMATS(4.1.0) software was used to analysis the splicing event. We used the *psiPerEvent* operation of SUPPA to calculate the Ψ values from the transcript quantifications obtained for all the alternative splicing events generated as described above with the *generateEvents* module of SUPPA (Alamancos GP, 2014). Data were visualised with R (R Core Team, 2017).

### Data and materials availability

All data needed to evaluate the conclusions in the paper are present in the paper and/or the Supplementary Materials. The RNA-Seq data from wild type and transgenic SMA mice is available for download from Gene Expression Omnibus (GEO) with accession number GSE206400.

### Statistics

ANOVA tests were used to compare the means between two and three or more groups, respectively. Statistics of survival times of SMA mice were determined by Kaplan-Meier estimation, and comparisons were made with the log-rank test. A two-way ANOVA was conducted to compare the effect of the treatment on weights of the animals using treatment as a between-subjects factor and time as a within-subjects factor. Power analysis was performed using G*Power 3.1.9.2 software (Erdfelder et al. 2009). GraphPad Prism version 8 was used to perform the statistical analyses (GraphPad, La Jolla, CA). A *P*-value less than 0.05 was set as statistically significant.

### Study approval

All animal procedures were authorised by the UK Home Office in accordance with the Animals (Scientific Procedures) Act 1986 and by the University of Oxford ethics committee (PPL no: PDFEDC6F0).

## Author contributions

AJK, SMH, MJAW, and CR conceived the study. AJK, NA, MH, AB, JS, and WFL performed the experiments. AJK and CR analysed the data and wrote the manuscript. All authors contributed to the discussion and writing of the manuscript.

## Acknowledgments

This study was supported by a Career Development Fellowship grant from the Wellcome Trust to CR (205162/Z/16/Z). AJK was supported by the Medical Research Council programme grant (BRT00040). WFL was supported by the Medical Research Council programme grant to M.J.A.W. (MR/N024850/1). The authors would like to thank the Structural Genomics Consortium for providing the library of small molecules.

