## Supplementary Data for "Type I PRMT inhibitor MS023 promotes *SMN2* exon 7 inclusion and synergizes with nusinersen to rescue the phenotype of SMA mice"

Supplementary material

**PRMT inhibition increases SMN expression and synergises with nusinersen to ameliorate SMA mice phenotype**

| Small molecule | Target protein family | Specific Targets | Concentration used for screening (Fig. 1) |
| --- | --- | --- | --- |
| A-485 | Acetyltransferase | p300, CBP | 10 µM |
| GSK484 | Arginine deiminases | PAD-4 | 10 µM |
| BAY-299 | Bromodomains | BRD1, TAF1 | 10 µM |
| BAY-850 | Bromodomains | ATAD2 | 2 µM |
| BAZ2-ICR | Bromodomains | BAZ2A, BAZ2B | 10 µM |
| BI-9564 | Bromodomains | BRD9, BRD7 | 10 µM |
| BSP | Bromodomains | pan-Bromodomain | 10 µM |
| GSK2801 | Bromodomains | BAZ2A, BAZ2B | 10 µM |
| GSK4027 | Bromodomains | PCAF, GCN5 | 10 µM |
| GSK6853 | Bromodomains | BRPF1 | 10 µM |
| GSK8814 | Bromodomains | ATAD2, ATAD2B | 10 µM |
| I-BRD9 | Bromodomains | BRD9 | 10 µM |
| I-CBP112 | Bromodomains | CREBBP, EP300 | 10 µM |
| JQ1(+) | Bromodomains | BRD2, BRD3, BRD4, BRDT (BET) | 10 µM |
| L-MOSES | Bromodomains | PCAF bromodomain | 10 µM |
| LP99 | Bromodomains | BRD9, BRD7 | 10 µM |
| NI-57 | Bromodomains | BRPF1, BRPF2, BRPF3 | 10 µM |
| NVS-CECR2-1 | Bromodomains | CECR2 | 1 µM |
| OF-1 | Bromodomains | BRPF1, BRPF2, BRPF3 | 10 µM |
| PFI-1 | Bromodomains | BRD2, BRD3, BRD4, BRDT (BET) | 10 µM |
| PFI-3 | Bromodomains | SMARCA,PB1 | 10 µM |
| PFI-4 | Bromodomains | BRPF1B | 10 µM |
| SGC-CBP30 | Bromodomains | CREBBP, EP300 | 10 µM |
| TP-472 | Bromodomains | BRD9, BRD7 | 10 µM |
| GSK864 | Dehydrogenase | Mutant IDH2 | 10 µM |
| GSK-J1 | Lysine Demethylase | JMJD3, UTX, JARID1B | 10 µM |
| GSK-LSD | Lysine Demethylase | LSD1 | 10 µM |
| T-26c | Matrix metalloproteinase | MMP-13 | 10 µM |
| A-395 | Methyl Lysine Binder | EED | 10 µM |
| UNC1215 | Methyl Lysine Binder | L3MBTL3 | 10 µM |
| A-196 | Methyltransferase | SUV420H1/H2 | 10 µM |
| A-366 | Methyltransferase | G9a, GLP | 10 µM |
| BAY-598 | Methyltransferase | SMYD2 | 10 µM |
| BAY-6035 | Methyltransferase | SMYD3 | 10 µM |
| GSK343 | Methyltransferase | EZH2 | 10 µM |
| GSK591 | Methyltransferase | PRTM5 | 10 µM |
| LLY-283 | Methyltransferase | PRMT5 | 10 µM |
| MS023 | Methyltransferase | Type I PRMTs | 10 µM |
| MS049 | Methyltransferase | PRMT4,6 | 10 µM |
| PFI-2 | Methyltransferase | SETD7 | 10 µM |
| PFI-5 | Methyltransferase | SMDY2 | 5 µM |
| SGC0946 | Methyltransferase | DOT1L | 10 µM |
| SGC3027 | Methyltransferase | PRTM7 | 5 µM |
| SGC707 | Methyltransferase | PRMT3 | 10 µM |
| TP-064 | Methyltransferase | PRMT4 | 10 µM |
| UNC0638 | Methyltransferase | G9a, GLP | 2 µM |
| UNC0642 | Methyltransferase | EHMT2 (G9a), EHTM1 (GLP) | 2 µM |
| UNC1999 | Methyltransferase | EZH2 | 5 µM |
| IOX1 | 2-oxoglutarate oxygenase | pan-2-OG | 10 µM |
| IOX2 | 2-oxoglutarate oxygenase | PHD2 | 10 µM |
| BAY-678 | Serine Proteases | Neutrophil Elastase | 10 µM |
| NVS-PAK1-1 | Serine/threonine-protein kinase | PAK1 | 10 µM |
| BAY-876 | Solute carriers | SLC2A1 | 10 µM |
| OICR-9424 | WD40 | WDR5 | 10 µM |

| Supplementary Table 1. List of small molecule epigenetic modulators |
| --- |

| Gene name | Alternative splicing event |
| --- | --- |
| *Apc* | Skipped exon |
| *Brsk2* | Skipped exon |
| *Camk2d* | Intron retention |
| *Eml4* | Skipped exon |
| *Gsap* | Skipped exon |
| *Klf12* | Mutually exclusive exon |
| *Mgat1* | Skipped exon |
| *Mtmr1* | Skipped exon |
| *Nin* | Skipped exon |
| *Nrcam* | Skipped exon |
| *Nsmf* | Skipped exon |
| *Pfkp* | Skipped exon |
| *Usp40* | Skipped exon |
| *Zfp420* | Skipped exon |

Supplementary Table 3. Splice events only corrected by the combination of MS023 and nusinersen

| Primer name | Primer sequence (5ʹ – 3ʹ) |
| --- | --- |
| Full length SMN2 forward primer | GCT TTG GGA AGT ATG TTA ATT TCA |
| Full length SMN2 reverse primer | CTA TGC CAG CAT TTC TCC TTA ATT |
| Delta 7 SMN2 forward primer | ACT TAC TAT CAT GCT GGC TG |
| Delta 7 SMN2 reverse primer | CCA GCA TTT CCA TAT AAT AGC C |
| Total SMN2 forward primer | GCG ATG ATT CTG ACA TTT GG |
| Total SMN2 reverse primer | GGA AGC TGC AGT ATT CTT CT |
| mouse Gapdh forward primer | AAAGGGTCATCATCTCCGCC |
| mouse Gapdh reverse primer | ACTGTGGTCATGAGCCCTTC |
| mouse PolJ forward primer | ACC ACA CTC TGG GGA ACA TC |
| mouse PolJ reverse primer | CTC GCT GAT GAG GTC TGT GA |
| human GAPDH forward primer | ACA TCG CTC AGA CAC CAT |
| human GAPDH reverse primer | TGT AGT TGA GGT CAA TGA AGG G |

Supplementary Table 4. Sequences of primers used throughout the study

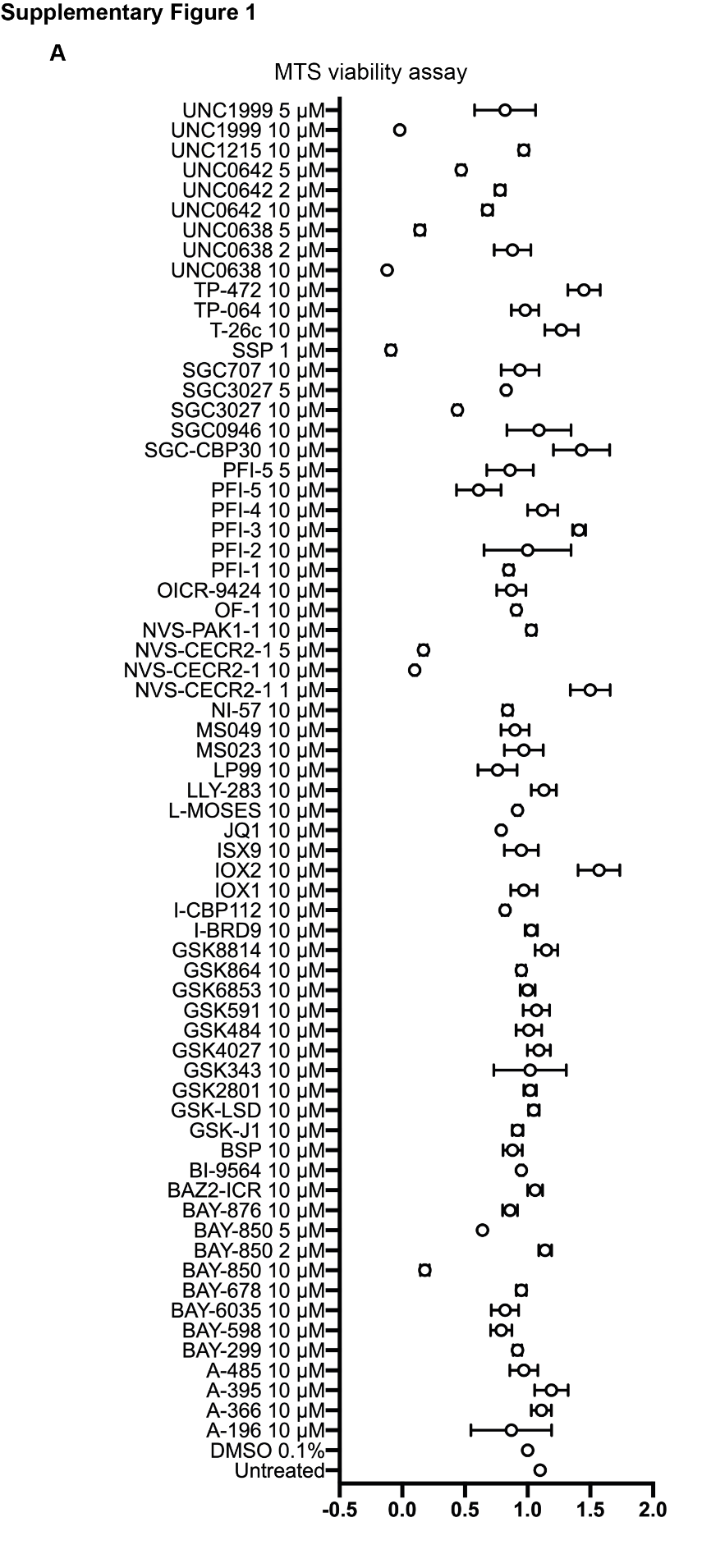

**Supplementary Figure 1. Effect of the epigenetic small molecules on SMA type II patient-derived fibroblasts viability. A**, Viability of cells, assayed by MTS assay, treated with epigenetic small molecules, relative to vehicle-treated cells (0.1% DMSO), normalised to one.

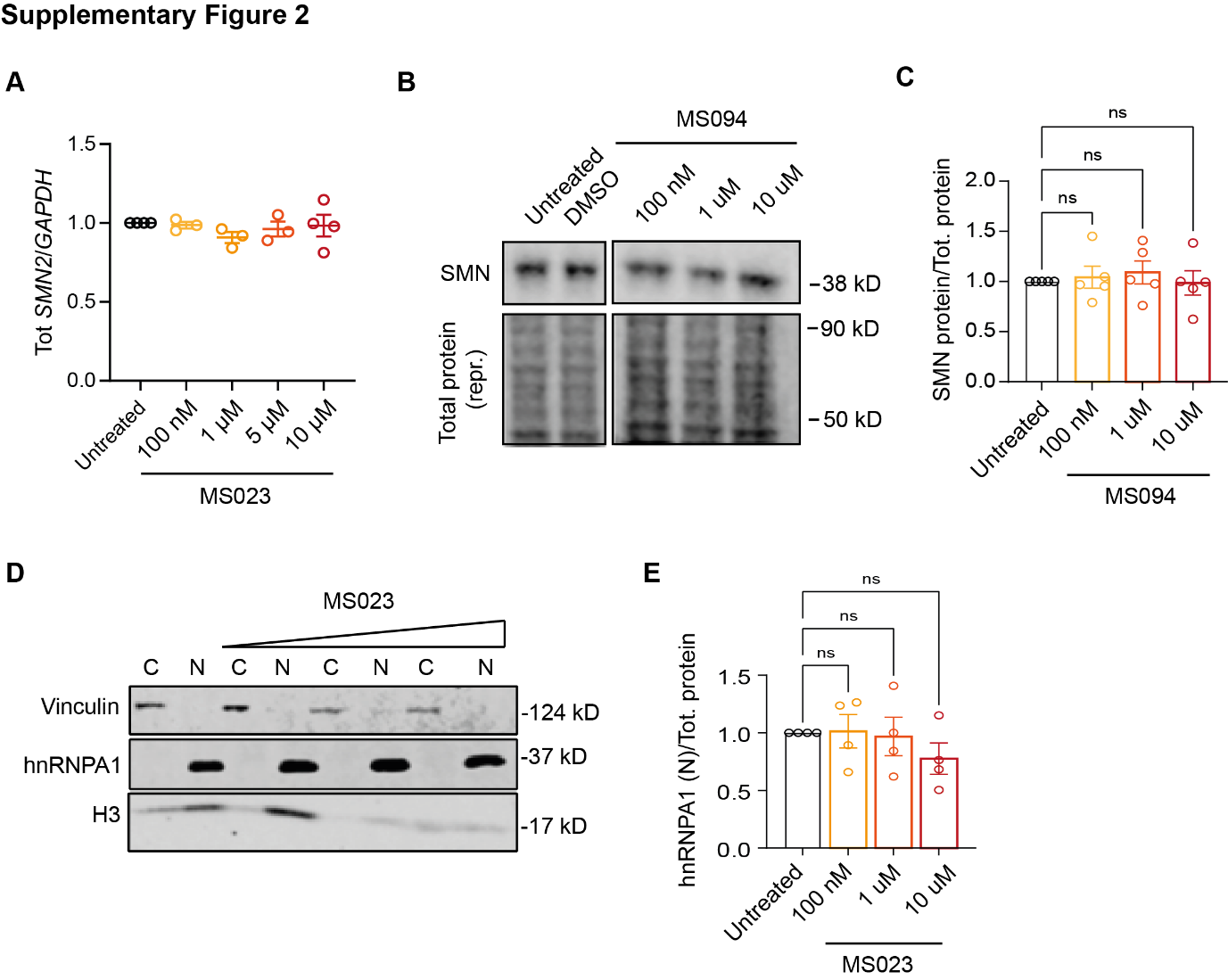

**Supplementary Figure 2.** **The increase in *SMN2* exon 7 inclusion by MS023 is specific and does not depend on altered hnRNPA1 levels. A**, SMA type II patient-derived fibroblasts were treated with the indicated concentration of MS023 (range: 100 nm–10 µM), (n = 3–4). Cells were harvested for RNA after 48 h incubation. Tot *SMN2* transcript levels relative to *GAPDH* are expressed as fold change compared to untreated SMA fibroblasts, normalised to one. Each dot represents a biological replicate (n = 3–4). **B**, Western blot showing SMN protein levels upon treatment with increasing MS094 (MS023 negative control) concentrations (top). A representative section of total protein stain, used for protein normalization, is shown (bottom). Size in kilodalton is indicated on the right. **C**, Quantification of SMN protein levels relative to total protein is shown. Each dot represents a biological replicate (*n* = 5). **D**, Western blot showing vinculin protein (top), hnRNPA1 (middle) and histone 3 (bottom) levels upon treatment with increasing MS023 concentrations and in cytoplasm (C) and nucleus (N). Size in kilodalton is indicated on the right. **E**, Quantification of hnRNPA1 protein levels relative to total protein is shown. Each dot represents a biological replicate (*n* = 4). **A**, **C** and **D**, Data are represented as mean ± s.e.m. and compared with a one-way ANOVA test with multiple comparisons.

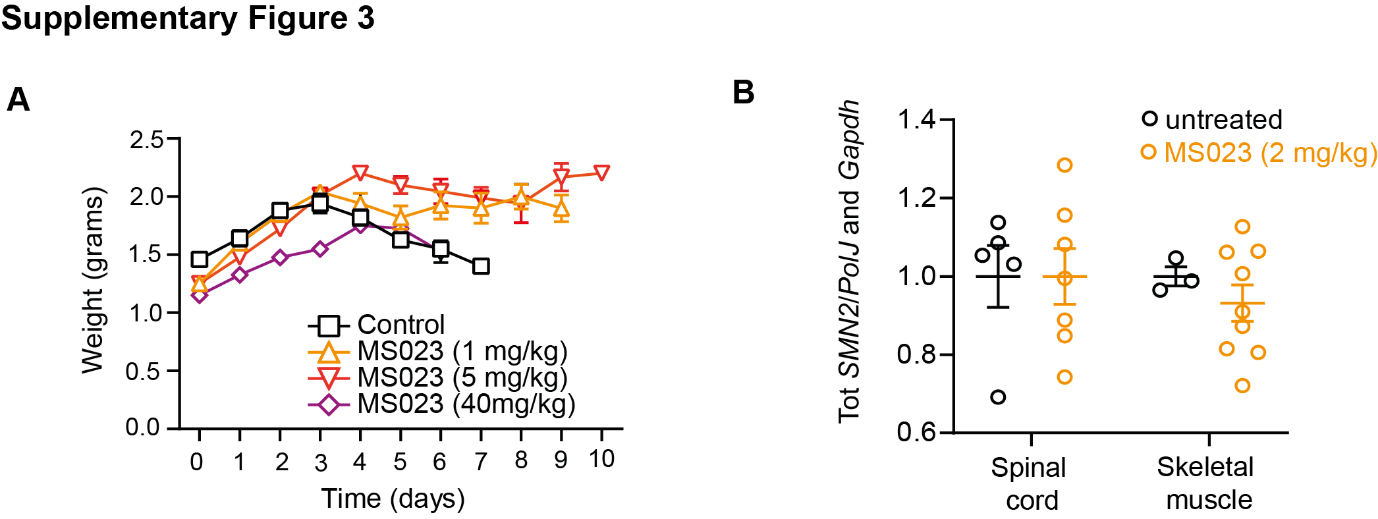

**Supplementary Figure 3. Oral administration of MS023 improves the phenotype of SMA mice. A**, Body weights of untreated, MS023- or vehicle-treated SMA mice from postnatal day 0 are shown. **B**, Tot *SMN2* transcript levels relative to *PolJ* and *Gapdh* in spinal cord and skeletal muscle of treated SMA mice compared to vehicle-treated SMA mice, normalised to one. Each dot represents a biological replicate (*n* = 10–12). **A**, **B**, Data are represented as mean ± s.e.m. and compared with a one-way ANOVA test with multiple comparisons.

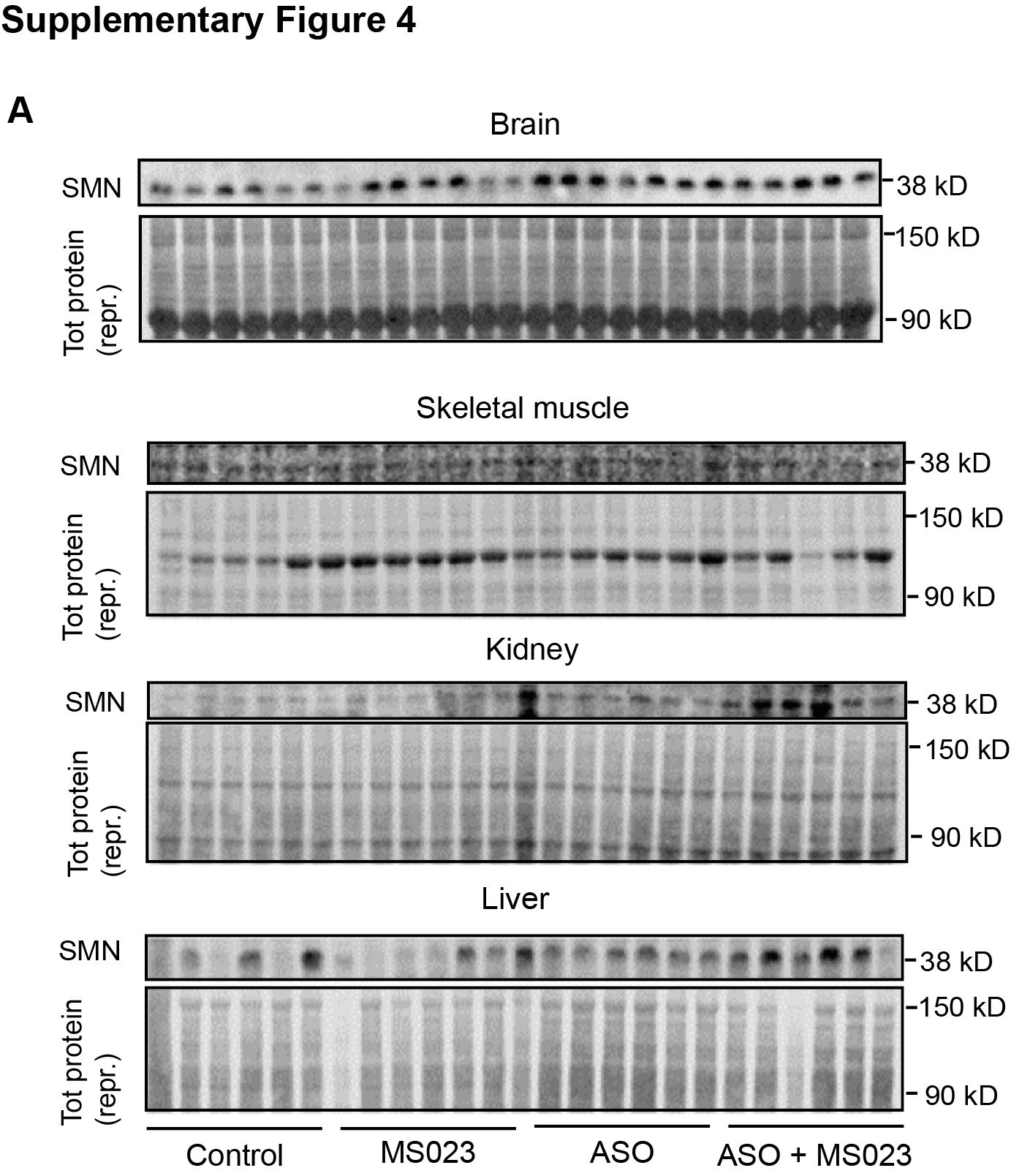

**Supplementary Figure 4. Combinatorial treatment with MS023 and ASO exerts synergistic effects in SMA mice. A**, Western blot showing SMN protein levels following the indicated treatments in brain, skeletal muscle, kidney and liver of SMA mice (top). A representative section of total protein stain, used for protein normalization, is also shown (bottom). Size in kilodalton is indicated on the right.

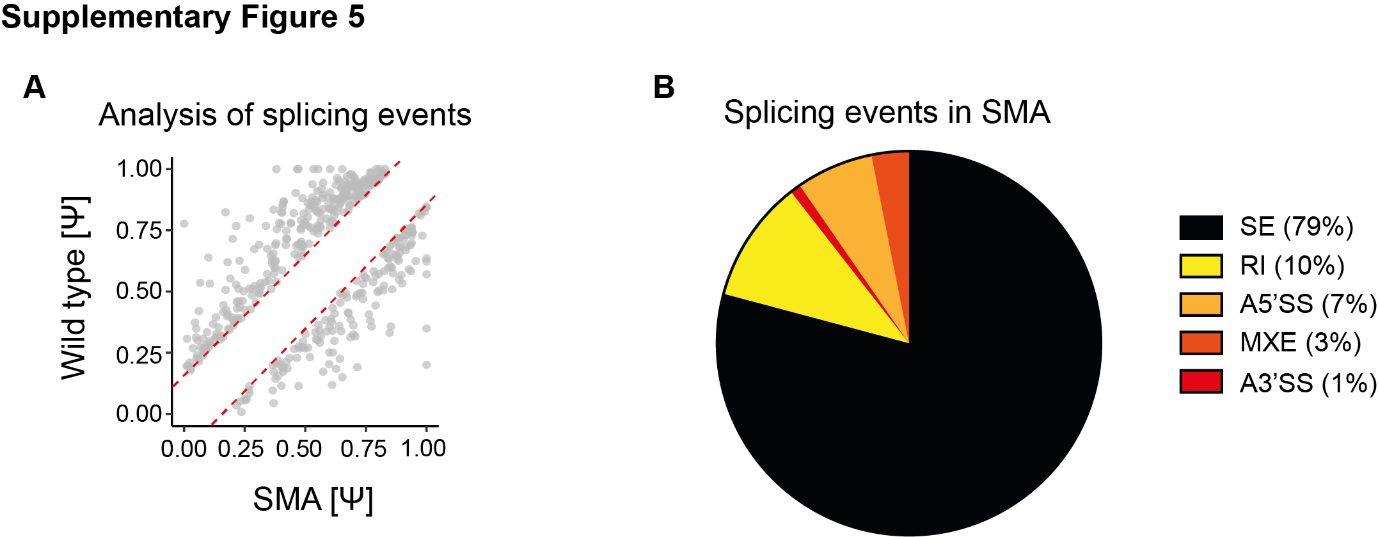

**Supplementary Figure 5. Altered splicing events in SMA. A**, Plot charts show the distribution of 446 aberrant splicing events (Ψ) in SMA mice relative to wild type littermates. The red dotted lines mark the ± 15% normalization range. **B**, Pie chart showing the proportion of altered splicing events in the spinal cord of SMA mice compared to wild type (SE: skipped exon, RI: intron retention, A5’SS: alternative 5’ splice site, MXE: mutually exclusive exon, A3’SS: alternative 3’ splice site).
